# Cocoa extract exerts sex-specific anti-diabetic effects in an aggressive type-2 diabetes model: a pilot study

**DOI:** 10.1101/2022.04.27.489764

**Authors:** Kathryn C. Racine, Lisard Iglesias-Carres, Jacob A. Herring, Mario G. Ferruzzi, Colin D. Kay, Jeffery S. Tessem, Andrew P. Neilson

## Abstract

Type 2 diabetes (T2D) is characterized by hyperglycemia and insulin resistance. Cocoa may slow T2D development and progression. This study employed male and female BTBR.Cg-*Lep*^*ob/ob*^/WiscJ (*ob/ob*) and wild type (WT) controls to assess the potential for cocoa to ameliorate progressive T2D and compare responses between sexes. Mice received diet without (WT, *ob/ob*) or with cocoa extract (*ob/ob* + c) for 10 weeks. Acute cocoa reduced fasting hyperglycemia in females, but not males, after 2 weeks. Chronic cocoa supplementation (6-10 weeks) ameliorated hyperinsulinemia in males and worsened hyperlipidemia and hyperinsulinemia in females, yet also preserved and enhanced beta cell survival in females. The underlying mechanisms of these differences warrant further study. If sex differences are apparent in subsequent preclinical studies, clinical studies will be warranted to establish whether these differences are relevant in humans. Sex differences may need to be considered when designing human dietary interventions for T2D.

## 1. Introduction

Diabetes is a global health crisis. The International Diabetes Federation reported in 2019 that ∼463 million adults worldwide were living with diabetes [1]. It is estimated that 90% of these are type 2 (T2D) [1]. While beta cell dysfunction is traditionally viewed as a late-stage T2D event, growing data suggest it may be an early event [2–4]. Obesity is a major risk factor for T2D, as it can result in insulin resistance [5,6]. During insulin resistance, perceived insulin demand is heightened, resulting in stress on beta cells-ultimately increasing beta cell mass and disrupting the balance of proliferation and apoptosis [7]. This combination of hyperinsulinemia and dysregulated beta cell physiology leads to beta cell exhaustion and failure and ultimately hypoinsulinemia in late T2D [8].

Cocoa (*Theobroma cacao*) is one of the most concentrated dietary sources of flavanols [9– 11]. Preclinical studies suggest a potential anti-diabetic role for cocoa flavanols [8]. Few studies have examined the impact of cocoa flavanols on beta cell function. Pancreatic beta cells secrete insulin and are responsible for maintenance of glucose homeostasis, along with glucagon-secreting alpha cells. Physiological conditions accompanying T2D, such as oxidative stress and inflammation, eventually induce beta cell failure and death and accelerate disease progression [5]. Promising studies suggest that flavanols protect beta cells *in vitro* [7,12–14] and *in vivo* [15,16], but this has not been investigated as thoroughly as effects in adipose tissue and skeletal muscle. We previously demonstrated that flavanols enhance beta cell function by increasing mitochondrial respiration [17]. Human studies of cocoa in the context of obesity and hyperglycemia have had mixed results [18,19]. Sex has been a largely ignored biological variable out of experimental convenience, however the potential differences due to sex have complicated translational research from male rodents to female humans. To harness the benefits of cocoa in humans, more information from preclinical models is needed regarding the role of disease state and sex in determining cocoa flavanol efficacy.

Rodent T2D models include insulin resistance and beta cell failure phenotypes. Monogenic models include *ob/ob* and *db/db* mice that are deficient in leptin signaling, consuming excess food and suppressing energy expenditure [20,21]. Environmental models include high fat feeding and/or streptozotocin (STZ) administration. High fat feeding involves slow development of beta cell damage, while STZ rapidly induces hyperglycemia by damaging beta cells. We employed an aggressive T2D model with the potential to reach beta cell damage. To achieve this, the *ob/ob* mutation on the BTBR (black and tan, brachyuric) background (BTBR.Cg-*Lep*^*ob*^/WiscJ) was used with wild type (WT) controls [22–24]. BTBR *ob/ob* mice progress rapidly to late-stage T2D and experience a loss of specific microRNAs in pancreatic islets, potentially resulting in an inability to increase beta cell replication in response to obesity [25].

Our objective was to perform a pilot study to identify sex differences in the response of T2D mice to cocoa supplementation during aggressive T2D.

## 2. Materials and Methods

### 2.1 Reagents

Standards of (−)-epicatechin, (±)-catechin, and procyanidin B2 were obtained from ChromaDex (Irvine, CA). Folin-Ciocalteu reagent and 4-dimethylaminocinnamaldehyde were obtained from Sigma-Aldrich (St. Louis, MO).

### 2.2 Cocoa flavanol-rich extract production and characterization

Flavanol-rich cocoa extract (CE) was prepared and analyzed by the Folin-Ciocalteu and 4-dimethylaminocinnamaldehyde methods [26,27], thiolysis [26,28] and UPLC-MS/MS [29]. Methodological details are found in Supplementary Material.

### 2.3 Animals

Approval was obtained from the Institutional Animal Care and Use Committee at the David H. Murdock Research Institute (#20-011), **and animal experiments followed** all policies in the *Guide for the Care and Use of Laboratory Animals*. Four-week-old BTBR.Cg-*Lep*^*ob*^/WiscJ mice (stock # 004824) were obtained from Jackson (Bar Harbor, ME): *N*=12 males and 12 females homozygous for the *Lep*^*ob*^ mutation (*ob/ob*), and *N*=6 male and 6 female wild-type (WT) controls. Mice were housed under standard conditions and allowed access to food and water *ad libitum* except where specified.

### 2.4 Diets

Mice were fed the 10% fat, 7% sucrose control diet from the Diet-Induced Obesity series (D12450J, Research Diets, New Brunswick, NJ) alone or +0.8% CE (Supplementary Table 1). This dose was based on estimated 10% CE yield from cocoa powder and studies[30,31] demonstrating that 8% cocoa powder in the diet blunts diet-induced obesity without altering food intake (details in Supplementary Information). Based on body surface area [32], this corresponds to 65 mg CE/kg/day in adult humans. For a 60 kg individual, this is 3900 mg CE or 39 g cocoa powder/d, equivalent to 8 doses of cocoa powder (5 g/dose), or 2/3 of an 80 g chocolate bar with 70% cacao. Treatments: WT, *ob/ob, ob/ob* + cocoa extract (*ob/*ob + c) (*n*=3/sex/group).

### 2.5 Body weight and food intake

Body weight and food intake were measured weekly.

### 2.6 Blood glucose

Fasting blood glucose was measured from the tail vein at weeks 2 and 6 after a 4 h fast using OneTouch Ultra Blue test strips (LifeScan, Inc., Milpitas, CA) and a glucometer (upper limit was 600 mg/dL; values above that were noted and recorded as 600 mg/dL).

### 2.7 Euthanasia

Animals were fasted 12 h and euthanized by CO_2_, followed by cervical dislocation. Blood was collected via cardiac puncture and liver and pancreata were preserved in 10% neutral buffered formalin (Thermo) overnight and then 70% ethanol. Blood was collected in serum tubes, clotted at room temperature (30 min) and serum prepared by centrifuging (2000 × *g*, 10 min). Samples were stored at −80°C.

### 2.8 Blood biomarkers

Fasting insulin and triglycerides from blood collected at euthanasia were quantified using a mouse ultrasensitive insulin ELISA kit (Crystal Chem, Elk Grove Village, IL) and a Triglyceride Quantification Colorimetric/Fluorometric Kit (Sigma).

### 2.9 Histology

Histological analysis of pancreata was conducted per established methods, methodological details can be found in Supplementary Information.

### 2.10 Genotyping

Formalin fixed liver tissue was used to genotype each animal to confirm the presence or absence of the *ob/ob* mutation. Samples were sent to Transnetyx (Cordova, TN) and results can be found in Supplementary Information.

### 2.11 Statistics

Data were analyzed using GraphPad Prism v9.1.2 (La Jolla, CA). When values were missing, a mixed-effects model was used. When initial one-way or two-way ANOVA indicated a significant overall treatment effect or interaction (P<0.05), individual means were compared using Tukey’s or Sidak’s *post-hoc* tests to control for multiple comparisons (α = 0.05).

## 3. Results

### 3.1 Cocoa

The composition of cocoa extract is shown in Supplementary Table 2. Yield of CE from powder was 18%.

### 3.2 Genotyping

We excluded one male (*ob/ob* + c) that did not carry the *ob* mutation (Supplementary Table 3).

### 3.3 Body weight

No statistically significant differences were observed in weight gain (Figure 1), likely due to significant variability in *ob/ob* mice on control diet. This could be due to weight loss or lack of weight gain (despite *ob/ob-*induced hyperphagia) often exhibited in severe diabetes [33]. WT mice had similar % weight gain compared to *ob/ob* controls, due to higher initial weights of *ob/ob* mice (Supplementary Figure 1). *Ob/ob* mice ate more food than WT controls (Supplementary Figure 1B, D). CE did not reduce food intake, and observed effects of CE were not due to reduced caloric intake [30,31]. The average CE intake was ∼1450 mg kg/d (Supplementary Figure 2).

**Figure 1.**
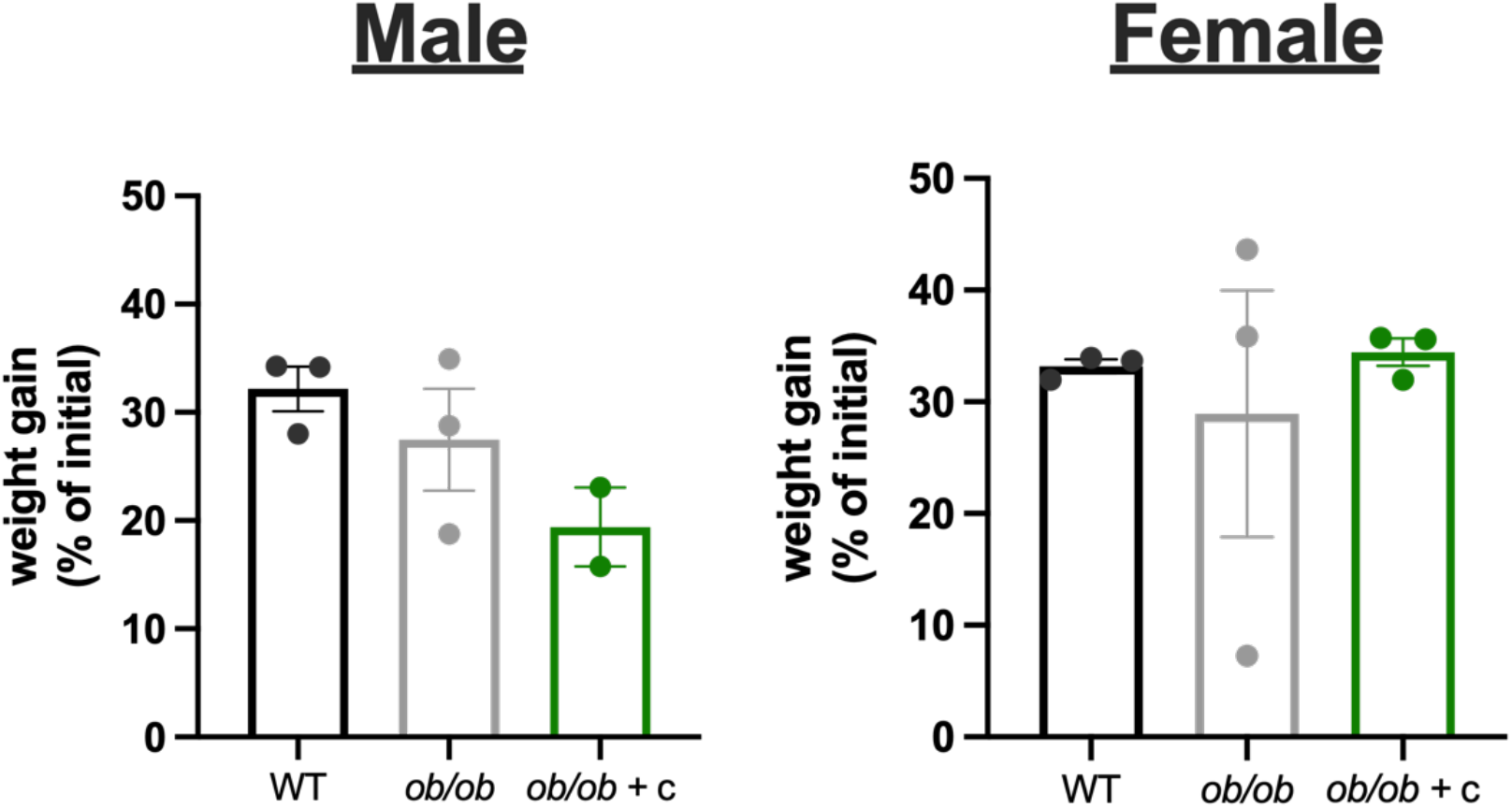
Weight gain (% of initial, mean ± SEM). Data were analyzed by 1-way ANOVA; due to lack of observed overall treatment effects, no *post hoc* tests were performed.

### 3.4 Fasting glucose

For males, fasting glucose was elevated in *ob/ob* and *ob/ob* + c mice compared to WT controls at 2 and 6 weeks (Figure 2). In females, distinct effects were observed. At week 2, *ob/ob* females had significantly elevated fasting glucose compared to WT controls, but supplementation with CE prevented this increase. At week 6, fasting blood glucose had worsened for both *ob/ob* and *ob/ob* + c, and the protective effect of CE was lost.

**Figure 2.**
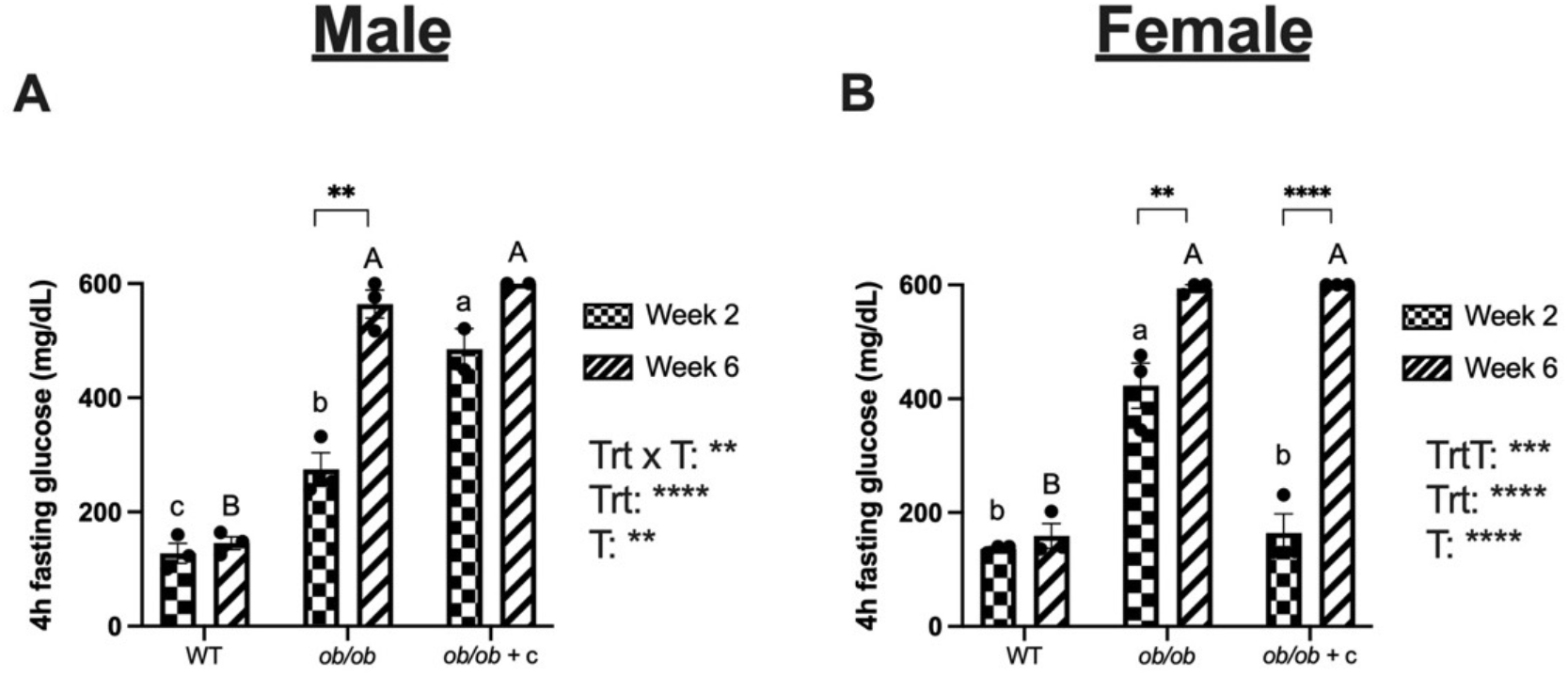
Four hour fasting blood glucose for males and females (mean ± SEM). Data were analyzed by 2-way ANOVA: treatment (Trt), time (T), and significance of main effects/interactions shown. If a significant main effect or interaction was detected, Sidak (time) or Tukey’s (treatment) *post hoc* test for multiple comparisons were performed: *, **, *** and **** indicate P ≤ 0.05, P ≤ 0.01, P ≤ 0.001 and P ≤ 0.0001, respectively. Lower-case and upper-case superscripts indicate significance for week 2 and week 6, respectively.

### 3.5 Insulin and triglycerides

Fasting blood insulin levels are shown in Figure 3A-B. In males, fasting insulin was elevated in *ob/ob* compared to WT controls. This was reversed by CE. In females, fasting insulin was elevated in ob/ob compared to WT controls, and CE further increased levels. Triglyceride levels are shown in Figure 3C-D. In males, no effect of treatment on serum triglycerides was detected. In females, serum triglycerides were not elevated in ob/ob mice compared to WT controls (as in males), but CE resulted in significant increases.

**Figure 3.**
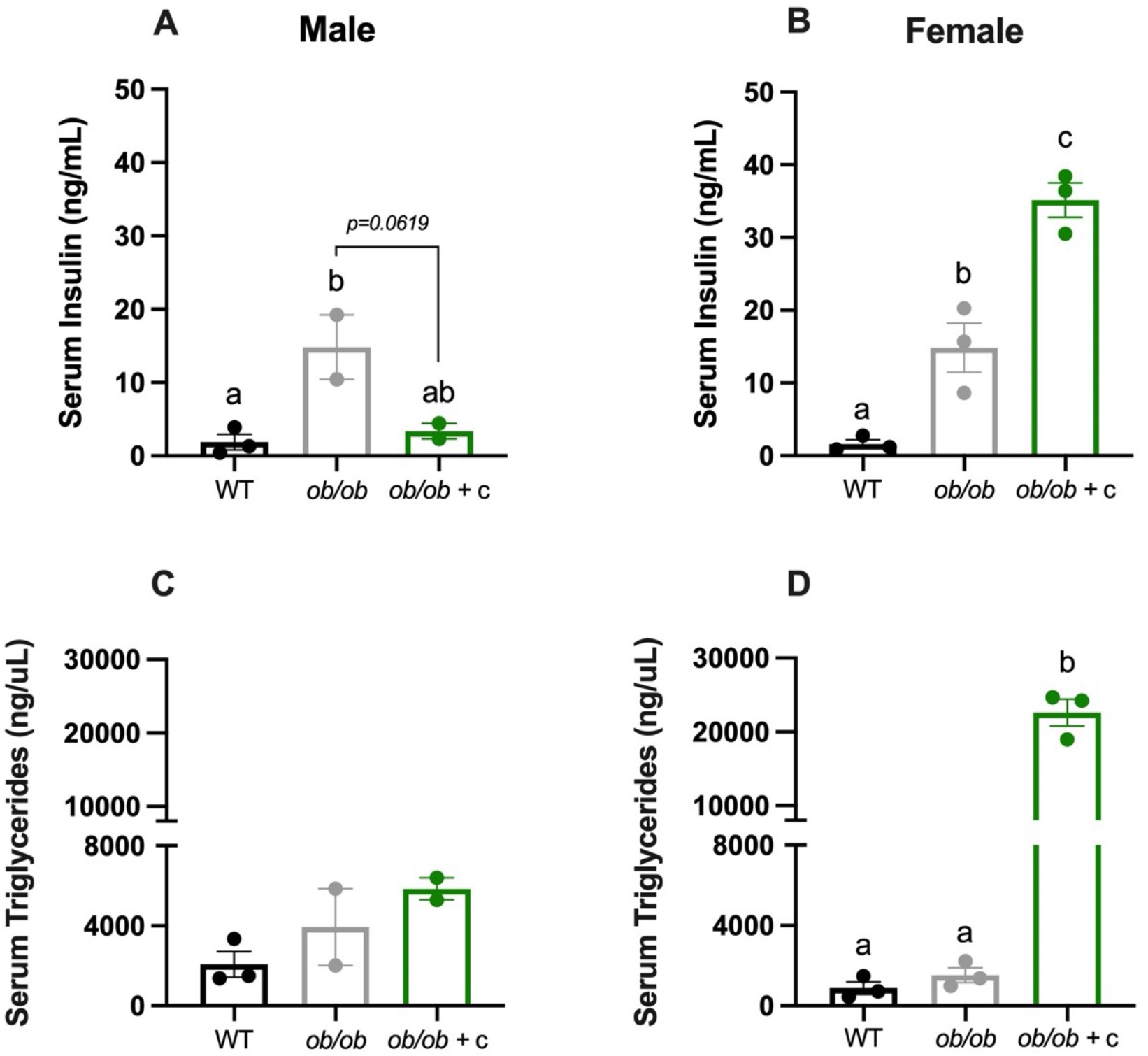
Fasting blood insulin (A, B, mean ± SEM) and serum triglycerides (C, D, mean ± SEM). Data were analyzed by 1-way ANOVA. If a significant treatment effect was detected, Tukey’s *post hoc* test was performed. Bars not sharing a common superscript are significantly different (P<0.05).

### 3.6 Pancreatic beta cell area

The % insulin positive area was quantified to determine beta cell survival (Figure 4). In males, the overall treatment effect was only borderline significant (*p=*0.073), and the trend suggested that WT had slightly greater beta cell area overall compared to *ob/ob* and *ob/ob* + c treatments. In females, *ob/ob* controls had significantly reduced beta cell area compared to WT, but CE protected against beta cell loss.

**Figure 4.**
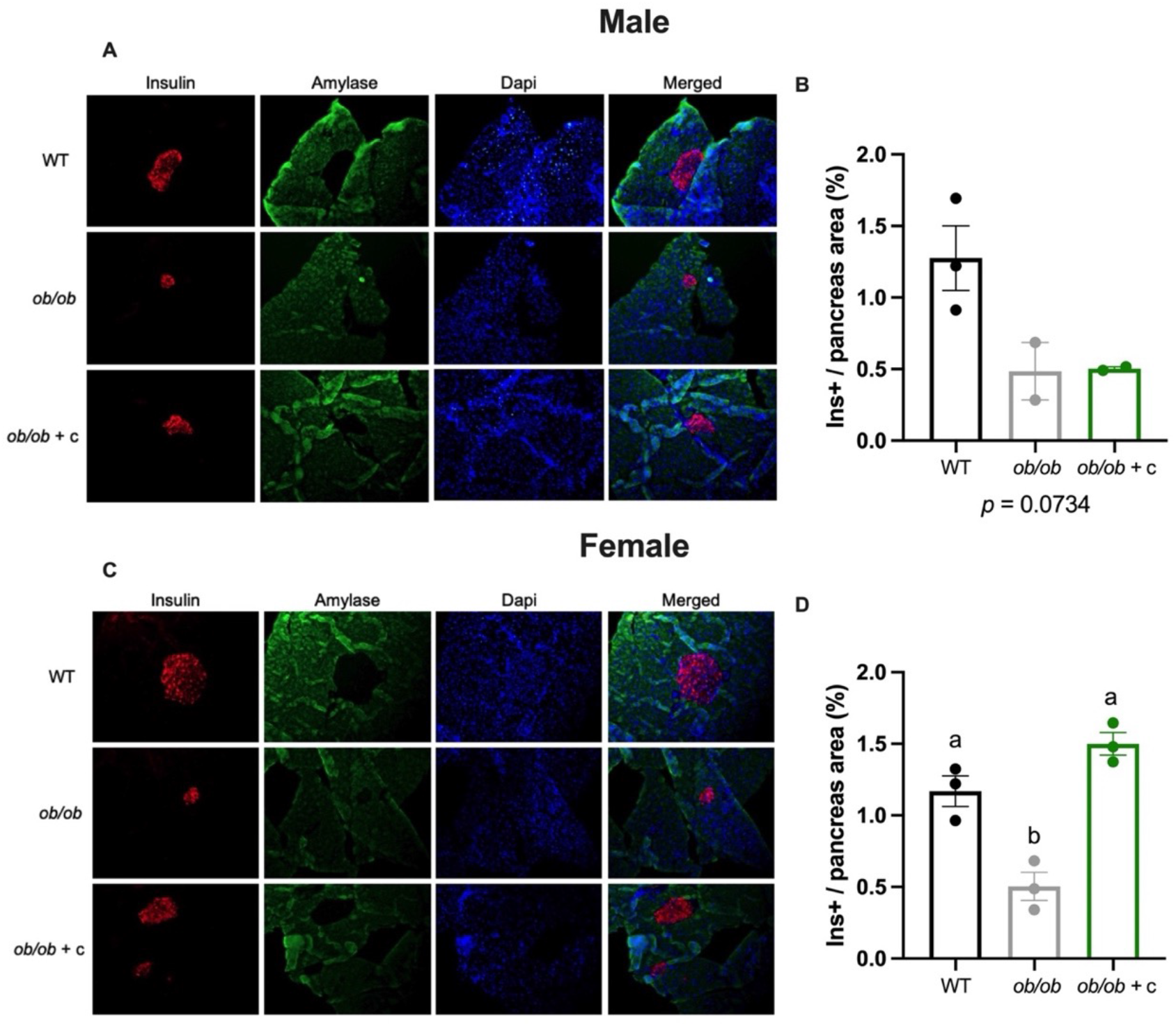
Representative pancreata images (males: A, females: C). Total % insulin positive area, (males: B, females: D, mean ± SEM). Images were captured at 20x magnification and analyzed using cellSens and ImageJ software for five section slides/animal. Data were analyzed by 1-way ANOVA. If a significant treatment effect was detected, Tukey’s *post hoc* test was performed. Bars not sharing common superscripts are significantly different (P<0.05).

## 4. Discussion

We compared effects of cocoa in male and female BTBR.Cg-*Lep*^*ob*^/WiscJ mice, an aggressive hyperglycemia model. This is the first study of flavanols *ob/ob* T2D mice on the BTBR background, and one of few studies to compare the effects of cocoa on both sexes [8,34]. Despite the small sample size, our results suggest significant differences between male and female *ob/ob* mice when fed a diet supplemented with CE.

Sex differences in weight gain were unclear due to lack of statistical significance facilitated by a small sample size. The speed of T2D progression was notable, as hyperglycemia and hyperinsulinemia are accelerated with the *ob* mutation on the BTBR background compared to B6 *ob/ob* [6,22]. Cocoa exerted protective effects early in the intervention in females. *Ob/ob* + c females exhibited significant protection against fasting hyperglycemia compared to *ob/ob* control at week 2, but this was lost by week 6 (Figure 2). The data further suggest that cocoa effects reverse based on sex as T2D progresses. Cocoa exerted a borderline significant protective effect on fasting hyperinsulinemia in males but worsened it in females (Figure 3A-B). Cocoa protected against beta cell loss in females (Figure 4) and together with fasting insulin expression, these results potentially indicate that in females, CE protected beta cell health and function even though it did not provide relief against fasting hyperglycemia. Conversely, CE reduced hyperinsulinemia in males, but that protection was not carried across to beta cells. We previously reported that cocoa monomers and cocoa flavanol microbial metabolites promote beta cell stability and enhance/stimulate beta cell function *in vitro* [17,35]. These effects appear to translate *in vivo* in female mice, but it remains unknown as to the mechanisms behind these effects. If these effects are translatable to humans, cocoa may be useful at different stages of beta cell function and insulin resistance based on sex.

CE did not affect fasting serum triglycerides in males but caused large increases in females, with significantly higher levels than *ob/ob* control (Figure 3C-D). The mechanism for this, as well as its implications, are unknown as these differences were not reflected in body weights (Figure 1). The combination of elevated circulating insulin and serum triglycerides, particularly in female *ob/ob* + c, can potentially be explained by more severe hyperglycemia and hyperinsulinemia, which may have led to greater lipolysis in order to meet fuel demands. A recent study reported worsening glucose metabolism in a mice fed purified polyphenols diet-induced obese mouse model [36]. These data suggest that the potential anti-obesity and anti-diabetic benefits of cocoa may differ in mice based on T2D stage, biological sex, and/or hormonal status. Whether these sex-specific effects are seen in humans remains to be established. Future studies should also capture metabolic responses in liver, skeletal muscle, and adipose tissue to better understand these relationships between sex, beta cell function, and glucose homeostatic perturbation.

This was a pilot study, with the limitation of small sample size due to mouse cost and availability. Despite this, sex-specific effects were evident. Additionally, diet formulation was based on WT mice, thus the actual dose of cocoa consumed was elevated due to *ob/ob* hyperphagia. Future research is needed using more translatable doses. Given the preliminary findings, larger studies with more statistical power are warranted.

These findings highlight the need to factor in sex-specific effects when studying cocoa and diabetes in animals and humans [37,38]. Often, animal studies of a single strain and/or sex are utilized to study dietary bioactives in chronic disease [39,40]. Studies should be performed to elucidate whether sex differences are due to hormonal differences [41] and/or genetic background. We previously reported that the anti-diabetic activity of quercetin was dependent on genetic background in mice [37]. It is unknown whether the sex-specific effects of cocoa are limited to *ob/ob* mice or the BTBR background. These findings need to be evaluated in multiple preclinical mouse models of T2D. If the results are generalizable across strains, human work is warranted. If the results are specific to narrow genetic contexts, studies can be performed to identify the responsible genetic loci [42].

In conclusion, cocoa supplementation exerts sex-specific effects on T2D. Cocoa reduced hyperglycemia in females, but not males, in the short term. Chronic supplementation ameliorated hyperinsulinemia in males and worsen hyperlipidemia and hyperinsulinemia in females, yet also preserve and enhance beta cell survival in females. The underlying mechanisms of these differences warrant further study. These pilot data will inform experiments with larger sample sizes. If sex differences are apparent in subsequent preclinical studies, clinical studies will be warranted to establish whether these differences are relevant in humans. Sex differences may need to be considered when designing human interventions for T2D.

## Acknowledgements

The authors acknowledge Dr. Glicerio Ignacio, DVM and Daniel Peralta for contributions to animal care and husbandry, Lauren Essenmacher for assistance with animal husbandry and blood glucose testing, and Lyric Ramsue for assistance with ELISAs.

