## Supplementary Information for "Cocoa extract exerts sex-specific anti-diabetic effects in an aggressive type-2 diabetes model: a pilot study"

---

### MATERIALS AND METHODS

*Cocoa flavanol-rich extract production and characterization* [1–5].

*Extract production.* Briefly, commercially available non-alkalized natural cocoa powder (The Hershey Co., Hershey, PA) was defatted three times by dispersion in hexane (1:3.75 cocoa:hexane), incubated for 10 min with sonication, centrifugation (5 min, 5000 × *g*) and discarding the supernatant. Residual hexane was evaporated at room temperature. Flavanols were extracted from defatted cocoa by dispersion in extraction solution (70:28:2 acetone:water:acetic acid, v/v/v; 1:3.75 defatted cocoa:extraction solution), incubated for 10 min with sonication, centrifugation (5 min, 5000 × *g*) and collecting the supernatant. This process was repeated three times, supernatants were pooled, and solvent evaporated under rotary evaporation (40-45 °C). The remaining CE was freeze dried and stored at –80°C.

*Folin-Ciocalteu colorimetric assay.* Cocoa extract (*n=3*) was diluted with 40% EtOH to a final concentration of 0.2 mg/mL. A standard curve of gallic acid was prepared at 0.0-1.0 mg/mL. Each sample and standard was diluted 10X with $ddH_2O$, followed by addition of 2N Folin-Ciocalteu reagent (1:2.5 sample solution:Folin). Sodium carbonate solution (7.5%, v/v) was added to all samples and standards. Samples and standards were incubated for 2 h at room temperature and then read at 765 nm. Total polyphenol concentration is expressed as mg Gallic Acid Equivalents (GAE).

*4-dimethylaminocinnamaldehyde (DMAC) colorimetric assay.* DMAC solution was prepared by combining stock HCl with EtOH (1:10 HCL:EtOH) and chilling at 4°C for 15 min. DMAC powder was added to the chilled solution at 0.001 g/mL. Cocoa extracts were diluted with EtOH to a final concentration of 100 ppm. A standard curve of procyanidin B2

(PCB2) was prepared at 0-100 ppm. DMAC solution was added to each sample and standard (1:5 sample:DMAC), mixed thoroughly, and read at 640 nm. Total flavanol concentration is expressed as mg PCB2 equivalents.

*Thiolysis.* Cocoa extract was diluted with MeOH to 0.5 mg/mL and then mixed (50 µL) with 50 µL HCl (3.3%, water) and 100 µL benzyl mercaptan (5%, MeOH). Samples were placed in a 90°C water bath for 5 min and then cooled on ice for 5 min. Unthiolyzed controls were prepared with cocoa extract and MeOH without heating in the water bath. Each thiolyzed sample (100 µL) was combined with 900 µL of 0.1% formic acid in water and 0.1% formic acid in ACN (95:5 v/v). Samples were analyzed on a Waters Acquity H-Class separations module with an Acquity UPLC HSS T3 column (2.1 mm × 100 mm, 1.8 µm) at 40°C. Binary gradient elution was performed using 0.1% formic acid in water (Phase A) and 0.1% formic acid in ACN (Phase B). Solvent flow rate was 0.6 mL/min and the linear gradient elution was as followed: 95% A (0-0.5 min), 65% A (6.5 min), 20% A (7.5-8.6 min), 95% A (8.7-10.5). (−)-electrospray ionization (ESI) together with tandem mass spectrometry (MS/MS) was used to analyze UPLC effluent on a Waters Acquity triple quadrupole (TQD) MS. (−) mode electrospray ionization (ESI) was performed with capillary, cone, and extractor voltages of −4.24 kV, 30.0 V, and 3.0 V respectively. Source temperature was 150°C and desolvation temperature was 400°C. Cone gas flows at a rate of 75 L/h and desolvation gas at 900 L/h. Argon (0.25 mL/min) was used as the collision gas in MS/MS. Multi-reaction monitoring (MRM) with a mass span of 0.2 Da was performed on parent ions and collision-induced dissociation (CID) on daughter ions. Inter-channel delays and interscan time was 1.0 s each. Additional calculations are done to account for the native monomers and are reported as DP of total flavanols. mDP oligomers and polymers and mDP of total flavanols are calculated as follows:

$$mDP\ (O+P) = \frac{\text{net number of monomers} + \text{net number of thiolytic derivatives}}{\text{net number of monomers}}$$

$$mDP\ (total\ flavanols) = \frac{\text{total monomers} + \text{net number of thiolytic derivatives}}{\text{total monomers}}$$

*UPLC-MS/MS.* A Waters Acquity H-class separation module equipped with a Waters Acquity UPLC HSS T3 column (2.1 mm × 100 mm, 1.8 µm, 40°C) and VanGuard HHS T3 precolumn (1.8 µm). Samples were maintained at 10°C. Binary gradient elution was performed with 0.1% (v/v) aqueous formic acid (phase A) and 0.1% (v/v) formic acid in acetonitrile (phase B). Solvent flow rate was 0.6 mL/min and the linear gradient elution was carried out as followed: 95% A (0-0.5 min), 65% A (6.5 min), 20% A (7.5-8.75 min), 95% A (8.85-10 min). An injection volume of 10 µL was used for all samples and standards. MS/MS analysis of column effluent was performed by (−)-ESI on a Waters Acquity TQD mass spectrometer equipped with a Z-spray electrospray interface. Ionization settings are as follows: −4.25 kV ESI capillary voltage, 150°C source temperature, 400°C desolvation temperature. $N_2$ was used for cone and desolvation gasses with flow rates of 75 L/h and 900 L/h, respectively. Ar was used as a collision gas. A standard curve of (–)-epicatechin, catechin, procyanidin B2, procyanidin C1, and cinnamtannin A2 (CinA2) was prepared and flavanol concentrations (procyanidin tetramer – decamer) are expressed as CinA2 equivalents.

*Diets.* Based on an estimated extraction yield of 10% from cocoa powder and food intake of ~0.1 kg diet/kg body weight/day in WT mice, the 0.8% dose was designed to provide ~800 mg CE/kg body weight/day (equivalent to ~8000 mg cocoa powder/kg body weight/day) to mice eating normal amounts of food (as we had not previously worked with *ob/ob* or BTBR mice). Based on body surface area conversion [6], this dose corresponds to 65 mg CE/kg body weight/day (~650 mg cocoa powder/kg body weight/day) in adult humans. For a 60 kg individual, this would equate to 3900 mg CE or 39 g cocoa powder. This equates to roughly 8 doses of cocoa powder (5 g/dose). The treatment groups for each sex were as follows: WT, *ob/ob*, *ob/ob* + cocoa extract (*ob/*ob + c) (*n*=3/sex/group). Diet treatments were started on the first day of week 1.

*Histology*. Pancreata were embedded in paraffin and five sections were cut from each animal. Sections were deparaffinized in xylene and rehydrated using a graded ethanol series. Antigen retrieval was performed using a sodium citrate buffer (Vector, H-3300-250) according to manufacturer's protocol. For insulin staining, slides were incubated overnight with guinea pig anti-insulin antibody (Fitzgerald, 20-IP35) followed by detection with an AlexaFluor 555-conjugated (red) goat-anti guinea pig secondary antibody (ThermoFisher, A-21435). For total tissue section area, slides were incubated overnight with rabbit α-Amylase (Sigma, A8273), followed by detection with an AlexaFluor 488-conjugated (green) goat anti-rabbit secondary antibody (ThermoFisher, A-32731). Slides were counterstained with DAPI. Images were captured at 20x magnification and analyzed using cellSens and ImageJ software for five tissue section slides per animal. β-cell area was determined as the total insulin-stained area divided by the total pancreatic tissue area (amylase and insulin-stained area) per slide.

### TABLES AND FIGURES

**Supplementary Table 1. Mouse diet formulations**

| | **Control diet** | | **Cocoa extract diet** | |
|---|---|---|---|---|
| **Macronutrient** | **% (g)** | **% (kcal)** | **% (g)** | **% (kcal)** |
| Protein | 19 | 20 | 19 | 20 |
| Carbohydrate | 67 | 70 | 67 | 70 |
| Fat | 4 | 10 | 4 | 10 |
| Total (%) | | 100 | | 100 |
| kcal/g | 3.8 | | 3.8 | |
| **Ingredient** | **g** | **kcal** | **g** | **kcal** |
| Casein | 200 | 800 | 200 | 800 |
| L-Cystine | 3 | 12 | 3 | 12 |
| Corn Starch | 506.2 | 2025 | 506.2 | 2025 |
| Maltodextrin 10 | 125 | 500 | 125 | 500 |
| Sucrose | 68.8 | 275 | 68.8 | 275 |
| Cellulose, BW200 | 50 | 0 | 50 | 0 |
| Soybean Oil | 25 | 225 | 25 | 225 |
| Lard | 20 | 180 | 20 | 180 |
| Mineral Mix S10026 | 10 | 0 | 10 | 0 |
| Dicalcium Phosphate | 13 | 0 | 13 | 0 |
| Calcium Carbonate | 5.5 | 0 | 5.5 | 0 |
| Potassium Citrate, 1 H2O | 16.5 | 0 | 16.5 | 0 |
| Vitamin Mix V10001 | 10 | 40 | 10 | 40 |
| Choline Bitartrate | 2 | 0 | 2 | 0 |
| Cocoa Extract | 0 | 0 | 8.51 | 0 |
| FD&C Yellow Dye #5 | 0.04 | 0 | 0 | 0 |
| FD&C Red Dye #40 | 0 | 0 | 0.05 | 0 |
| FD&C Blue Dye #1 | 0.01 | 0 | 0 | 0 |
| **Total** | 1055.05 | 4057 | 1063.56 | 4057 |

**Supplementary Table 2. Cocoa extract characterization**

| Measure | Value |
|---|---|
| **Total polyphenols (Folin)**<br>(mg GAE/mg extract ± SEM[a]) | 0.25 ± 0.01 |
| **Total flavanols (DMAC)**<br>(mg PCB2/mg extract ± SEM[b]) | 0.20 ± 0.01 |
| **Mean degree of polymerization (thiolysis)**<br>(mDP ± SEM)[c] | Including monomers: 2.14 ± 0.02<br>Excluding monomers: 2.55 ± 0.05 |
| **Procyanidin characterization (LC-MS/MS)**<br>(mg/g extract ± SEM) | |
| Catechin | 8.56 ± 0.10 |
| Epicatechin | 18.9 ± 0.36 |
| Dimer | 5.69 ± 0.16 |
| Trimer | 6.74 ± 0.23 |
| Tetramer | 3.93 ± 0.043 |
| Pentamer | 3.61 ± 0.078 |
| Hexamer | 1.61 ± 0.11 |
| Heptamer | 1.27 ± 0.14 |
| Octamer | 0.664 ± 0.027 |
| Nonamer | 0.376 ± 0.041 |
| Decamer | 0.119 ± 0.019 |

[a]Gallic acid equivalents
[b]Procyanidin B2 equivalents
[c]Mean degree of polymerization (monomer residues per flavan-3-ol molecule)

**Supplementary Table 3. Mouse genotype results**

| Sample ID[a] | Detected *ob/ob* mutation |
|---|---|
| **Male** | |
| WT (A) | -- |
| WT (B) | -- |
| WT (C) | -- |
| *ob/ob* (A)[b] | NA |
| *ob/ob* (B) | ++ |
| *ob/ob* (C) | ++ |
| *ob/ob* + *c* (A) | ++ |
| *ob/ob* + *c* (B) | ++ |
| *ob/ob* + *c* (C) | -- |
| **Female** | |
| WT (A) | -- |
| WT (B) | -- |
| WT (C) | -- |
| *ob/ob* (A) | ++ |
| *ob/ob* (B) | ++ |
| *ob/ob* (C) | ++ |
| *ob/ob* + *c* (A) | ++ |
| *ob/ob* + *c* (B) | ++ |
| *ob/ob* + *c* (C) | ++ |

[a]A-C designate individual animals within each group

[b]unable to genotype, mouse died at week 10 prior to euthanasia

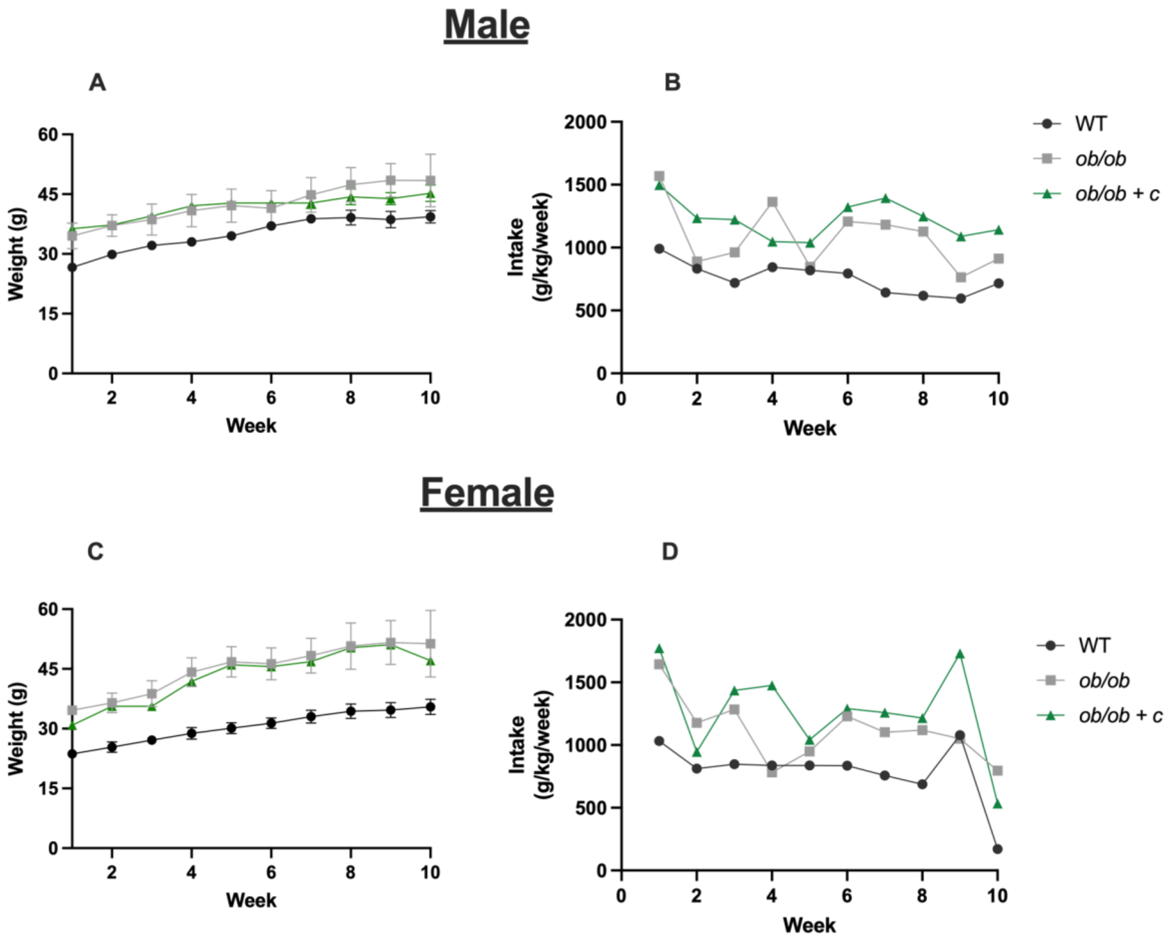


***Supplementary Figure 1.*** *Absolute weights during the 10-week experiment for male (A, B) and female (C, D) mice. Male mice: weight gain over time (A), total absolute weight gain (B). Female mice: weight gain over time (C), weight gain over time, total absolute weight gain (D. Dots represent individual animals; colored bars and error bars represent mean ± SEM. For bar graphs, data were analyzed by 1-way ANOVA; due to lack of observed overall treatment effect for any graph, no post hoc tests were performed to compare treatment means. Error bars and statistical analyses are not shown for food intake, as each value is from n=1 cage.*

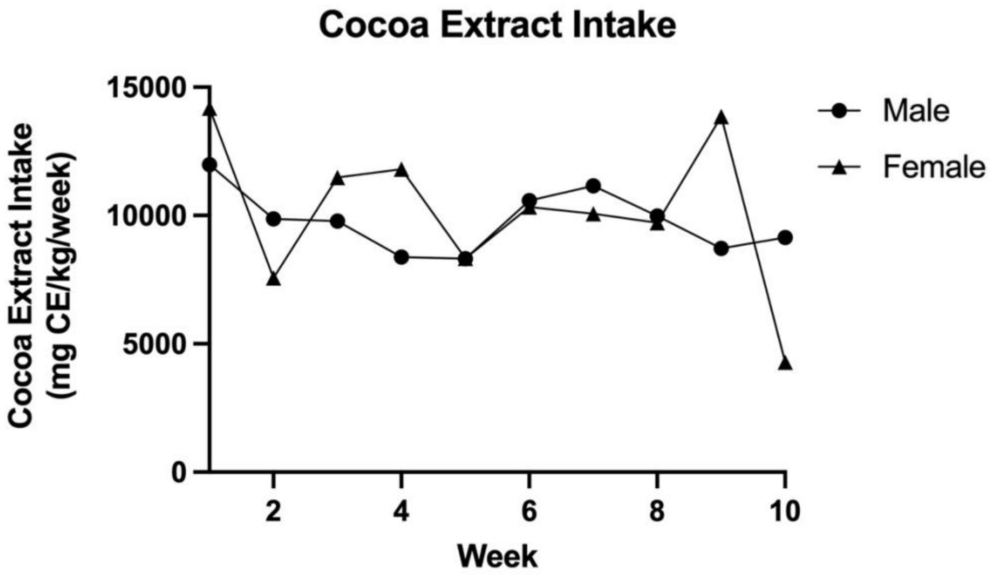


***Supplementary Figure 2.*** *Cocoa extract intake for both male and female mice. Male and female treatments without cocoa extract supplementation are not shown Error bars and statistical analyses are not shown for extract intake, as each value is from n=1 cage.*
